# A versatile platform strain for high-fidelity multiplex genome editing

**DOI:** 10.1101/410001

**Authors:** Robert G. Egbert, Harneet S. Rishi, Benjamin A. Adler, Dylan M. McCormick, Esteban Toro, Ryan T. Gill, Adam P. Arkin

## Abstract

Precision genome editing accelerates the discovery of the genetic determinants of phenotype and the engineering of novel behaviors in organisms. Advances in DNA synthesis and recombineering have enabled high-throughput engineering of genetic circuits and biosynthetic pathways via directed mutagenesis of bacterial chromosomes. However, the highest recombination efficiencies have to date been reported in persistent mutator strains, which suffer from reduced genomic fidelity. The absence of inducible transcriptional regulators in these strains also prevents concurrent control of genome engineering tools and engineered functions. Here, we introduce a new recombineering platform strain, BioDesignER, which incorporates (1) a refactored λ-Red recombination system that reduces toxicity and accelerates multi-cycle recombination, (2) genetic modifications that boost recombination efficiency, and (3) four independent inducible regulators to control engineered functions. These modifications resulted in single-cycle recombineering efficiencies of up to 25% with a seven-fold increase in recombineering fidelity compared to the widely used recombineering strain EcNR2. To facilitate genome engineering in BioDesignER, we have curated eight context-neutral genomic loci, termed Safe Sites, for stable gene expression and consistent recombination efficiency. BioDesignER is a platform to develop and optimize engineered cellular functions and can serve as a model to implement comparable recombination and regulatory systems in other bacteria.

## INTRODUCTION

The design-build-test (DBT) cycle is a common paradigm used in engineering disciplines. Within the context of synthetic biology it is employed to engineer user-defined cellular functions for applications such as metabolic engineering, biosensing, and therapeutics (1, 2). The rapid prototyping of engineered functions has been facilitated by advances in *in vitro* DNA assembly, and plasmids have traditionally been used to implement designs *in vivo* given their ease-of-assembly and portability. However, for deployment in contexts beyond the laboratory such as large-scale industrial bioprocesses or among complex microbial communities, plasmid-based circuits suffer from multiple limitations: high intercellular variation in gene expression, genetic instability from random partitioning of plasmids during cell division, and plasmid loss in environments for which antibiotic use could disrupt native microbial communities or is economically infeasible (3, 4). These shortcomings can be ameliorated once a design is transferred from a plasmid to the host genome, which offers improved genetic stability and lower expression variation (5) along with reduced metabolic load (6). However, behaviors optimized for plasmid contexts often do not map predictably to the genome. As such, building and testing designs directly on the genome can reduce the DBT cycle time and facilitate engineering cellular programs for complex environments.

Expanding synthetic biology efforts to genome-scale engineering has historically been limited by factors such as low endogenous rates of recombination, lack of optimized workflows for recombination, and uncertainty due to locus-dependent expression variability (7, 8). The advent of recombination-based genetic engineering (recombineering), which relies on homologous recombination proteins, often *exo, bet*, and *gam* from bacteriophage λ, in conjunction with linear donor DNA containing target homology and the desired mutations, has enabled genomic deletions, insertions, and point mutations at user-defined loci (9-13). Recombineering has enabled generation of genomic discovery resources such as the *E. coli* K-12 in-frame, single-gene deletion collection of non-essential genes (Keio collection) (14) and technologies such as trackable multiplex recombineering (TRMR), which enables genome-scale mapping of genetic modifications to traits of interest (15, 16). In addition, pooled library recombineering approaches such as CRISPR-enabled trackable genome engineering (CREATE) have combined CRISPR-Cas9 gene editing schemes with barcode tracking to enable high-throughput mutational profiling at single-nucleotide resolution on a genome-wide scale. (17).

Meanwhile, techniques such as multiplex automated genome engineering (MAGE) have been developed to generate complex mutagenesis libraries by extending recombineering to simultaneously modify multiple genetic loci through iterative cycles of single-strand DNA (ssDNA) oligonucleotide recombination (18). MAGE has enabled several genome-scale recombineering efforts such as the recoding of all 321 occurrences of TAG stop codons with synonymous TAA codons in a single *E. coli* strain (19, 20), the removal of all instances of 13 rare codons from 42 highly expressed essential genes to study genome design constraints (21), the insertion of multiple T7 promoters across 12 genomic operons to optimize metabolite production (22), and the His-tagging of 38 essential genes that encode the entire translation machinery over 110 MAGE cycles for subsequent *in vitro* enzyme studies (23). In addition, methods such as tracking combinatorial engineered libraries (TRACE) have been developed to facilitate the rapid, high-throughput mapping of multiplex engineered modifications from such genomic explorations to phenotypes of interest (24, 25).

To achieve the high levels of recombination necessary to carry out large-scale, multiplexed genome editing, many of these studies required the use of mutagenic strains. Specifically, the endogenous methyl-directed mismatch repair (MMR) system, which acts to revert newly made recombineering modifications when active, was removed to more effectively retain targeted modifications in the standard MAGE strain EcNR2. While deactivation of the MMR dramatically enhances recombination efficiency, it also increases the rate of background mutagenesis by 100-1000 fold (26, 27). Indeed, in converting all 321 occurrences of TAG stop codons to TAA stop codons, Lajoie *et al.* noted the addition of 355 unintended (i.e. off-target) mutations after the final strain construction (20).

Several approaches have been proposed to circumvent the use of MMR-deficient strains and thus avoid their high basal rates of off-target mutagenesis. Designs utilizing mismatches that are poorly repaired or that introduce silent mismatches near the desired mutation can be used to evade MMR, which only recognizes short mismatches (28). Furthermore, oligos containing chemically modified bases can be used to evade MMR correction and increase allelic-replacement efficiency (29). While these approaches boost recombination rates without increasing basal mutagenesis rates, they either limit the range of mutations that can be implemented or significantly increase oligonucleotide costs.

More recent efforts have focused on approaches to create a transient mutagenesis state. Specifically, cells are cycled between phases of elevated mutation rate, during which editing can take place efficiently, and phases of wild type-like mutation rates, during which cells can be propagated without incurring a significant number of background mutations. *Nyerges et al.* reported the use of a temperature-controlled mismatch repair deficient strain (*E. coli* tMMR) in which the MMR machinery can be transiently inactivated by shifting cells to a non-permissive temperature (36°C) during oligonucleotide incorporation and cell recovery and then reactivated by returning cells to the permissive temperature (32°C) for propagation (30). While this approach reduces the number of off-target mutations by 85%, it restricts cell growth to 32°C and hence increases the time between recombineering cycles. In contrast, *Lennen et al.* developed a plasmid-based MAGE system, Transient Mutator Multiplex Automated Genome Engineering (TM-MAGE). In TM-MAGE, *E. coli* Dam methylase is inducibly overexpressed to transiently limit MMR and thus enable high allelic replacement efficiencies with a 12- to 33-fold lower off-target mutation rate than strains with fully disabled MMR (31).

Given existing approaches to recombineering in *E. coli*, researchers still face a trade-off between high-efficiency genome editing and genome stability. Here we present a rational genome engineering approach to develop such a platform strain, called BioDesignER, with enhanced recombineering efficiency while retaining low off-target mutagenesis rates and enabling short editing cycle times. We refactored the λ-Red machinery in *E. coli* K-12 MG1655-derived EcNR1 to decrease cycle time and reduce toxicity, stacked genetic modifications shown to increase recombination rates, and characterized gene expression across the chromosome at curated integration loci, herein referred to as Safe Sites. We also introduced genomic modifications to independently control four transcriptional regulators of gene expression and characterized the induction regime for each regulator. We profiled the growth and ssDNA recombination rates of BioDesignER with a dual-fluorescent reporter cassette integrated at each Safe Site and also demonstrated the retention of double-strand DNA (dsDNA) recombination capabilities in the strain. We performed a comparative study of background mutagenesis rates of our strain and alternative platform strains using a fluorescent reporter-based fluctuation assay and found that BioDesignER exhibited a 4.2-fold lower mutagenesis rate compared to the widely used recombineering strain EcNR2. Finally, we compared the multi-cycle accumulation of targeted mutations for BioDesignER and other high-efficiency recombineering strains and found that BioDesignER exhibited similar multiplex editing efficiencies to EcNR2.nuc5-, a persistent mutator strain with the highest reported ssDNA recombination efficiency. BioDesignER is a high-fidelity genome engineering strain that uniquely enables high-efficiency recombineering while retaining low basal mutagenesis rates.

## MATERIALS AND METHODS

### Chemicals, reagents, and media

LB Lennox Medium (10 g/L Tryptone, 5 g/L Yeast Extract, 5 g/L NaCl; Sigma Aldrich, USA) was used to culture strains for experiments, to prepare electrocompetent cells for recombineering, and as recovery broth following electroporation. Antibiotics concentrations used were 34 μg/mL for chloramphenicol, 100 μg/mL for carbenicillin, and 50 μg/mL for kanamycin. Anhydrotetracycline (CAS 13803-65-1; Sigma Aldrich, USA) was used at 100 ng/mL to induce the λ-Red genes for recombineering. For *thyA*-mediated recombineering steps, M9 minimal media supplemented with 0.4% glucose, 0.2% casamino acids, thymine (100 μg/mL), and trimethoprim (50 μg/mL) was used. M9 minimal media with valine (20 μg/mL) was used to select for the *ilvG*^+^ genotype. All M9 minimal media was supplemented with biotin at 10 μg/mL to account for the biotin auxotrophy common to all EcNR1derivative strains.

### Oligonucleotides

Oligos were ordered from Integrated DNA Technologies (IDT), resuspended in 1x TE buffer at either 500 uM (recombineering oligos) or 100 uM (standard amplification oligos), and stored at −20°C. For recombineering workflows, oligos were designed to target the lagging strand of DNA replication and contain at least 35 bp of homology to the target locus. Oligos for testing recombination efficiency were ordered with 5’ phosphorothioate base modifications. Oligo sequences for individual lineage construction steps are available in linked Benchling files from Supplementary Table S1.

### Strains

Relevant strains used in this work are reported in Table 1. Supplementary Table S2 provides relevant genotypes of 32 intermediate strains in the lineage between EcNR1 to BioDesignER. This table includes descriptions of genetic modifications, associated recombineering, selection and enrichment methods, and a web link to sequence-level detail of each modification. Supplementary Table S3 provides a summary of strain identification numbers and genotypes for the BioDesignER lineage.

**Table 1.**
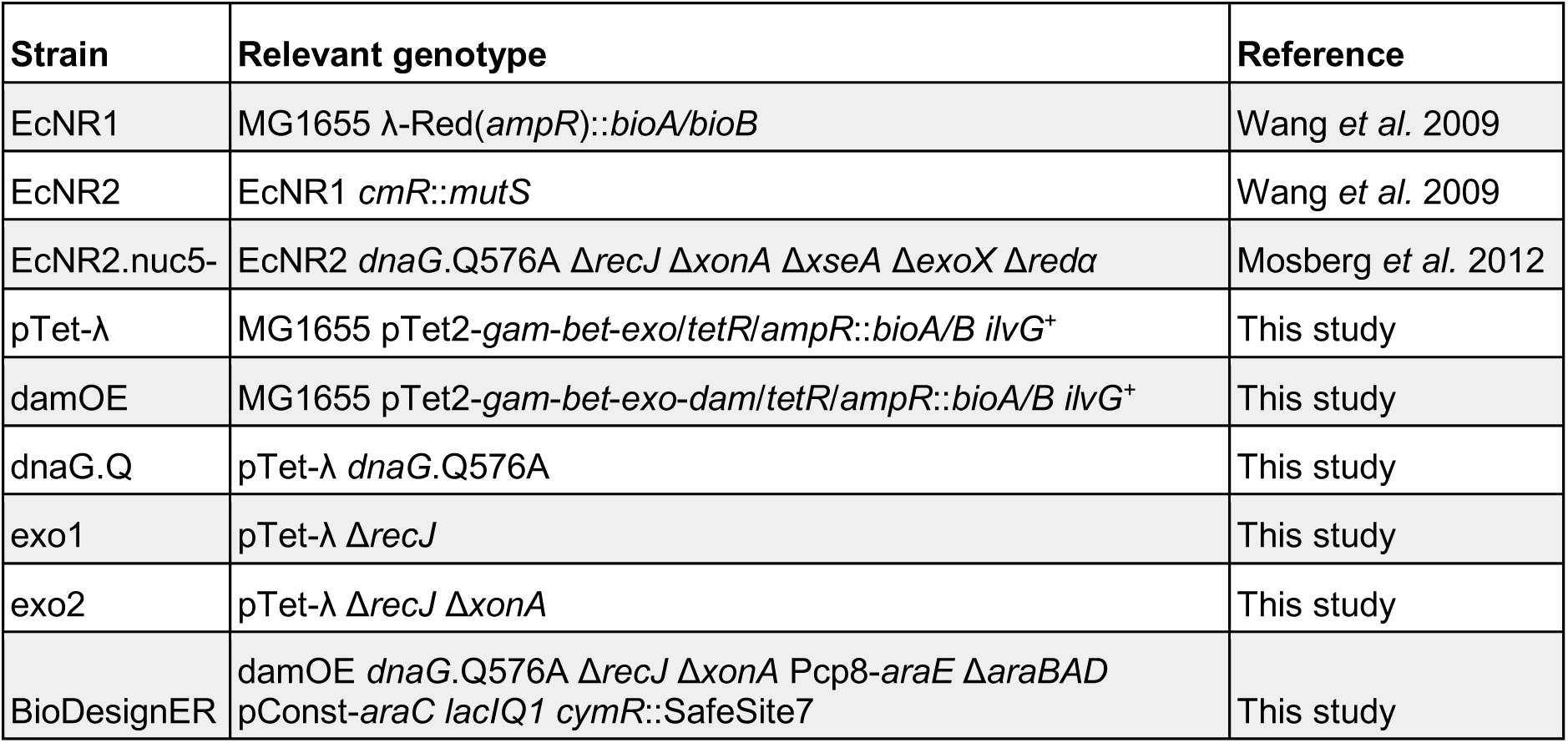
Genotypes of abbreviated strains.

### Growth rate measurements

Two clones of each strain were cultured overnight in LB Lennox (LB) medium with chloramphenicol. The following morning each strain culture was back-diluted 1:100 into two media types: (1) LB with aTc (LB+aTc) (2) LB. The resulting inocula were divided into four technical replicates and then grown for up to 18 hours in a Biotek Synergy 2 microplate reader. The growth rate at early exponential phase was calculated from the resulting optical density data using custom analysis scripts in python.

### Competent cell preparation and recombineering

Strains were grown overnight in LB Lennox medium (LB) with antibiotics as appropriate at 37°C. The following morning each strain culture was back-diluted 1:100 into 25 mL LB+aTc and grown at 37°C until they reached OD600 0.3-0.4. The resulting mid-log cultures were chilled in a 4°C ice-water bath. Cultures were centrifuged (Beckman-Coulter Allegra 25R) at 8000 xg and subjected to two washes: (1) 25 mL chilled water (2) 15 mL chilled 10% glycerol. The cell pellets after the final glycerol wash were resuspended in 10% glycerol, yielding approximately 500 uL of competent cells given the residual cell mass from the wash.

Due to their different induction and growth requirements, EcNR2 and EcNR2.nuc5- strains were grown overnight at 30°C, back-diluted 1:100 into 25 mL LB+chlor media, and cultured at 30°C until they reached OD600 0.3-0.6. The λ-Red machinery was induced by incubating the cultures in a 42°C water bath for 15 minutes after which the strains were chilled in a 4°C ice-water bath for at least 10 minutes. The remainder of the preparation for EcNR2 and EcNR2.nuc5- follows the same aforementioned wash steps.

40 uL of competent cells were used for each recombineering reaction. Oligos were diluted to 50 μM concentration in 10% glycerol and 10 μL of the diluted oligo was added to the competent cell mixture. For water control reactions, 10 μL of water was added. For multiplexed reactions, 10 μL of a cocktail with a total oligo concentration of 50 μM was used. The resulting cell-oligo mix was transferred to a chilled cuvette (1 mm gap, VWR) and electroporated using a BTX™-Harvard Apparatus ECM™ 630 Exponential Decay Wave Electroporator with the following parameters: voltage (1800 V), resistor (250 Ω), capacitor (25 μF).

### Fluorescence-coupled scar-free selection/counter-selection

Working from the *ΔthyA* strain RE095 and derivative strains (Supplementary Table S2), a dsDNA *thyA* cassette with or without a fluorescence gene (Supplementary Figure S1) was amplified with 35-50 bp homology to a target genomic locus and integrated via standard recombineering as described above, with the exception that cells were made competent by growing in LB supplemented with thymine (100 μg/mL) and trimethoprim (50 μg/mL). Integrants of *thyA* were selected for on LB media. Colonies with fluorescence-coupled *thyA* cassettes were screened visually for fluorescent phenotypes on a blue-light transilluminator. Proper insertion of the cassette was confirmed by locus-specific colony PCR. Replacement of the *thyA* cassette was performed through recombineering with a ssDNA or dsDNA cassette as described in Supplementary Table S2 and selected for on M9 agar plates supplemented with thymine (100 μg/mL), trimethoprim (50 μg/mL), and casamino acids (0.2%). Removal of fluorescence-coupled *thyA* cassettes was screened visually for non-fluorescent colonies via blue-light transillumination and sequences were validated via colony PCR and Sanger sequencing.

### Recombineering efficiency and Safe Site expression measurements

Competent cells were transformed with water (control) or oligos to turn off sfGFP, mKate2, or both reporters. Following electroporation, cells were resuspended in 3 mL LB+carb. These cultures were mixed and 30 μL was transferred into an additional 3 mL LB+carb for overnight growth at 37°C (or 30°C for EcNR2 and EcNR2.nuc5-). The following morning, saturated cultures of each transformation were diluted 1:200 into Phosphate-buffered saline (PBS) solution and run on a Sony SH800 cell sorter for single-cell flow cytometry analysis. At least 50,000 events were recorded for each reaction, and the fractional abundance of each reporter phenotype (GFP+ RFP+, GFP- RFP+, etc.) in the population was measured. The threshold for each reporter phenotype was determined via a prior calibration in which gates for each fluorescent reporter were measured. For measurement of gene expression across Safe Sites, overnight outgrowths of control reactions from the Safe Site recombineering efficiency transformations were processed on the flow cytometer.

### Response curves of inducible regulators

BioDesignER was transformed with plasmids containing a GFP gene regulated by each transcription factor - AraC (pBAD), CymR (pCym), or LacI (pLac) - and controlled by one of two replication origins, p15A or pSC101. Plasmids with the p15A origin contain a *kanR* marker and plasmids with the pSC101 origin use a *cmR* marker. Plasmid sequences are available via Benchling (https://benchling.com/organizations/arkinlab). Individual colonies were inoculated in LB with an appropriate antibiotic to maintain the plasmid and grown overnight. Saturated cultures were diluted 200fold into a microtiter plate (Corning 3904) and grown at 37°C with shaking in a Biotek H1 plate reader. Kinetic growth and fluorescence measurements were taken every 5 or 10 minutes for 12 hours. Absorbance was measured at 600 nm. GFP fluorescence was measured using 485/20 nm and 520/15 nm filter cubes for excitation and emission, respectively. mKate fluorescence was measured using 560/20 nm and 615/30 nm filter cubes for excitation and emission, respectively. Fluorescence values measured nearest OD 0.5 were used to estimate absorbance-normalized fluorescence in each channel.

### Flow cytometry analysis of inducible regulators

Saturated cultures from the kinetic growth assays used to measure regulator inducer responses were diluted 400-fold into PBS and analyzed in a BD LSR Fortessa flow cytometer (488 nm excitation / 525/50 nm emission for GFP; 561 nm excitation / 670/30 nm emission for mKate) using an autosampler. Raw .fcs files were imported for pre-processing and subsequent analysis with custom Python scripts (see Supplementary Data D1 for example) using the FlowCytometryTools software package (https://github.com/eyurtsev/FlowCytometryTools). For each sample, 50,000 events were captured and outliers in forward scatter and side scatter were removed using a filter with cut-offs for events outside the second and third quartile.

### Safe Site expression analysis

Data for expression levels at each Safe Site were calculated from flow cytometry data and used to measure recombineering efficiency at each Safe Site. Data files were extracted for four recombineering conditions (no oligo control, GFP-off, mKate-off, and dual-off) and two biological replicates. The geometric mean for each fluorescence channel was calculated from filtered data. Specifically, events outside the second and third quartiles for forward and side scatter channels were removed from analysis for each .fcs data file. The dual-fluorescent subpopulation for each measurement was extracted by gating at a value that excluded recombinant, non-fluorescent subpopulations but did not truncate the distribution of the dual-fluorescent subpopulation.

### Fluctuation assay

Fluctuation tests were performed on an inactivated *cmR-mNeon* translational fusion cassette integrated at Safe Site 1. The cassette was inserted using selection on chloramphenicol and subsequently inactivated for the following strains: pTet-λ, damOE, dnaG.Q + *ΔrecJ*/*ΔxonA*, BioDesignER, pTet-λ *ΔmutS*, and EcNR2. The inactivated cassette was first integrated as double-strand DNA into the respective strains via recombineering and selected for by plating on LB agar supplemented with 34 μg/mL chloramphenicol. A premature stop codon (AAA to TAA) was inserted into *cmR-mNeon* via single-strand DNA recombineering with an oligo harboring the stop codon mutation. The non-fluorescent population was enriched using cell sorting (Sony SH800) and the sorted cells were plated on LB agar plates.

Prior to fluctuation tests, individual non-fluorescent colonies were grown at 30°C in LB+carb and stored at −80°C as glycerol stocks normalized to OD600 of 0.5. For the fluctuation tests, cultures were diluted 1000-fold and grown for 16 hours in permissive conditions of LB+carb at 30°C (N=24). For pTet-λ *ΔmutS*, EcNR2, and EcNR2.nuc5-, 20μL of culture was spotted onto LB agar plates supplemented with chloramphenicol and carbenicillin. For all other strains, 100 μL volume spots were used. Viability counts were estimated for all strains by serial dilutions of 6 independent cultures on LB agar plates supplemented with carbenicillin. Chloramphenicol-resistant mutants were counted and mutation rates were inferred by the MSS-MLE method (32, 33).

### Iterative recombineering cycling

Strains were prepared for transformation using the competent cell protocol described above using 25 mL of culture with a target OD600 of 0.3. Each culture was resuspended in ~500 μL of 10% glycerol after washes. Each transformation consisted of 40 uL competent cells mixed with 10 μL of 50 μM oligo mix. After transformation, cells were recovered in 3 mL LB supplemented with carbenicillin. The recovery culture was grown to saturation before beginning the next round of competent cell prep and recombination. In parallel, the recovery culture was diluted 1:60 into an additional 3 mL of LB supplemented with carbenicillin and grown to saturation prior to measurements using flow cytometry (Sony SH800).

## RESULTS

### Rational strain design

We introduced multiple targeted modifications to an MG1655-derivative strain to decrease recombination cycle time, reduce toxicity of the recombination machinery, and introduce a transient hypermutation phenotype via hypermethylation (Figure 1A-B, Table 1). Using EcNR1 (18) as the host, we refactored the λ-Red recombination machinery, which consists of the genes *exo, bet*, and *gam*, and serves as the basis for mediating homology-directed recombination of ssDNA and dsDNA products. To reduce recombineering cycle times, we replaced the temperature-inducible regulation of the λ-Red locus with a TetR-regulated design (Figure 1C). This allowed us to propagate cells at 37°C instead of 30-32°C during all phases of a recombineering workflow: competent cell prep, λ induction, cell recovery, and selection. We also minimized the λ prophage by deleting the λ-*kil* gene, which has been reported to be responsible for the cell death phenotype observed under λ-Red expression (34), and other dispensable phage genes. Finally, we introduced DNA adenine methyltransferase (*dam*) to the λ-Red operon of our strain. Co-induction of *dam* with the λ-Red recombination genes results in transient hypermutation via hypermethylation, which has been reported to enable incorporated mutations to evade MMR (31).

**Figure 1.**
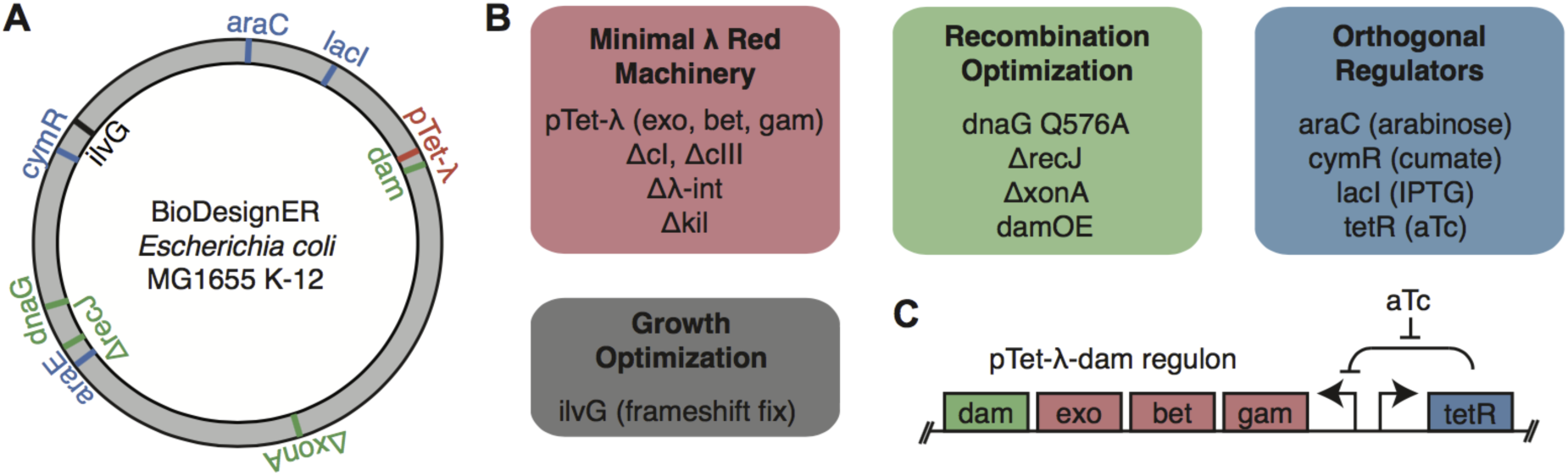
Overie of modification in BioDesignER. **(A)** Chrmosome map of the BioDesignER strain (derived fom *E coil* MG 1665K-12) with modification made in the platform strain mapped to corresponding positions on the genome. **(B)** Functional grouping of genomic modification based on purpose in platform strain (e.g. minimization of λ-Red machinery,optimization of recombination efficiency, implementation of multiple orthogonal reguators, or optimization of grpwth). **(C)** Genetic architecture of refactored λ-Red machinery and dam overex pression construct along with regulation by TetR.

To remove a valine-sensitive growth defect present in *E. coli* K-12, we restored expression of *ilvG*. K12 contains three acetohydroxy acid synthases (*ilvB, ilvG, ilvH*) that are involved in branch-chained amino acid biosynthesis. K-12 does not express *ilvG* due to a natural frameshift mutation and thus exhibits a growth defect in the presence of exogenous valine and the absence of isoleucine (35, 36). This valine-sensitive growth phenotype is alleviated by restoration of *ilvG* (37). Using oligo-mediated recombination (Methods) we removed the frameshift mutation in the endogenous *ilvG* gene, which has been reported to enable faster growth in minimal media. We called this strain pTet-λ.

We next incorporated genomic modifications shown to improve recombination efficiency. Using a scar-free genome engineering workflow that utilizes a novel *thyA* selection/counter-selection cassette containing a fluorescent marker (Supplementary Figure S1, Methods), we iteratively generated multiple beneficial mutations. For example, genetic variants of DNA primase (*dnaG*) enhance recombination efficiency by increasing the length of Okazaki fragments, thus exposing longer stretches of the lagging strand of the replication fork to ssDNA recombination (38). We incorporated the *dnaG*.Q576A variant, which was shown to boost recombination efficiency more than other *dnaG* mutants in EcNR2, into our strain.

Endogenous nucleases can degrade exogenous DNA used in recombineering workflows. The removal of a set of five nuclease genes (*endA, exoX, recJ, xonA, xseA*) has been shown to improve ssDNA recombination efficiency (39). However, while this exonuclease knockout strain, EcNR2.nuc5-, exhibited increased recombination efficiency, it also resulted in a lower post-electroporation growth rate compared to EcNR2. This suggested that deletion of the entire set of nucleases introduces to the strain an undesirable physiological defect. To avoid such growth defects, which are compounded for workflows requiring multiple recombineering cycles, we looked to systematically combine exonuclease knockouts that distinctly improve recombination rates. We constructed individual knockouts of each of the five exonucleases and measured the recombineering efficiency of the resulting strains. We assayed recombination efficiency for each exonuclease knockout using oligo mediated recombination at a genomically-encoded *sfGFP* reporter. In this assay, recombination of an oligo designed to introduce a premature stop codon into *sfGFP* results in a loss of fluorescence that can be quantified using flow cytometry. Deletions of *xonA* (4.2%) and *recJ* (2.6%) showed the greatest efficiencies, while the remaining exonuclease deletions yielded nominal efficiencies (<1%) (Supplementary Table S4). Based on these results, we deleted only two of the five exonucleases (*recJ, xonA*) in the next step of strain construction. While deletion of the λ-Red exonuclease (*exo*) can also promote stability of exogenous ssDNA (39), we opted to retain it due to its role in dsDNA recombination. The culmination of these genetic modifications in addition to the inducible regulator modifications described below, resulted in the BioDesignER strain.

To assess the effect of BioDesignER modifications on strain fitness, we measured the growth rates of key strains in the modification lineage in LB rich media (Figure 2A). We noted that, in general, doubling times decreased as additional modifications were made. Additionally, in contrast to cell death reported for extended co-expression of λ-*kil* with the recombination machinery (34), we observed only a slight increase in doubling time when expressing the refactored λ-Red cassette.

**Figure 2.**
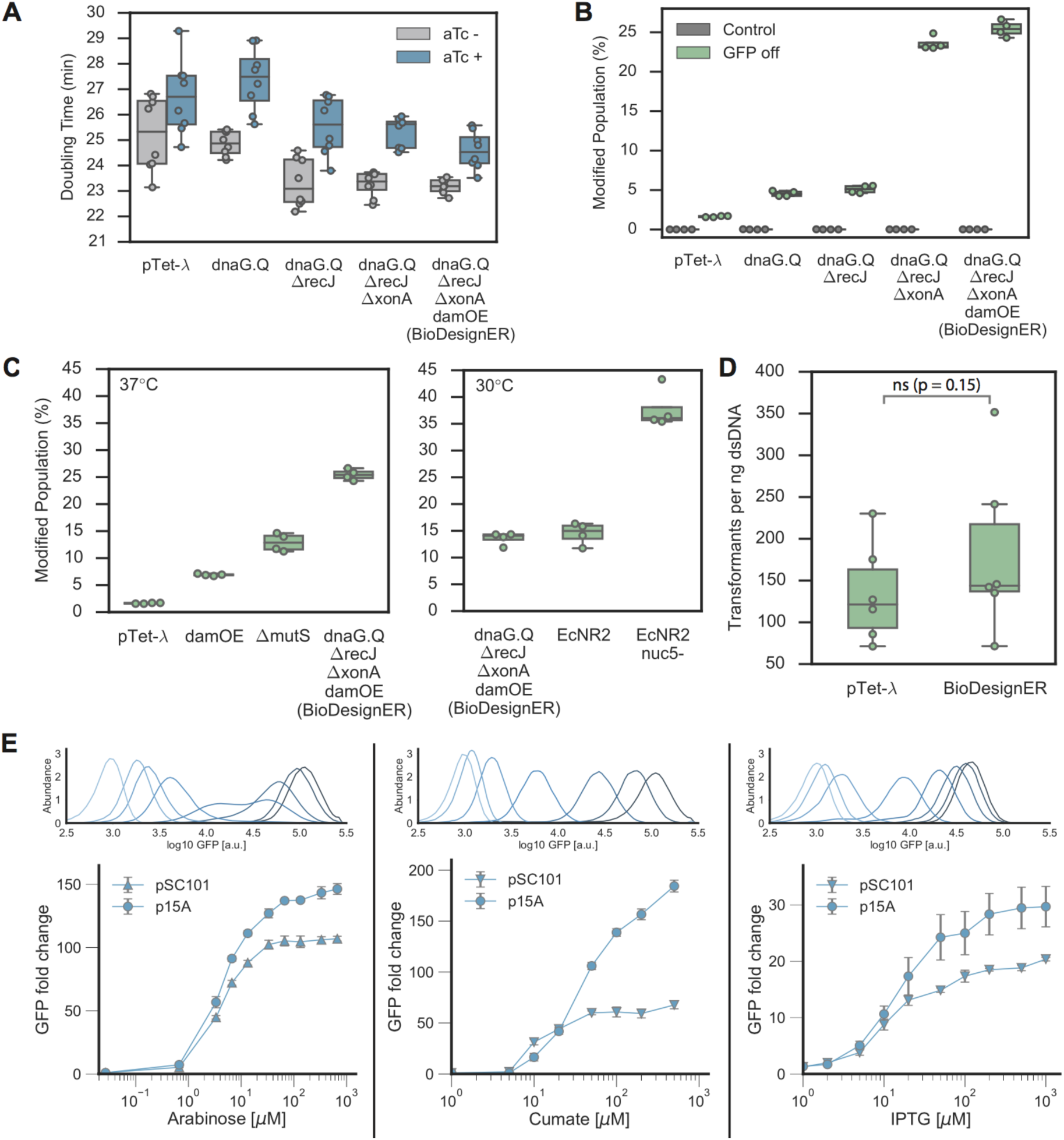
Strain Characterization. **(A)** Doubling times of strains growth at37°C for eleted strains of BioDesignsER lineage starting with the startin with Tet-λ Additional modifications shown moving to the right. Doubling times reported for strains grown with (blue) and without (gray) aTc induction to show the effect of λ-Red expression on growth. Data repersented as boxplots overlayed with corresponding data points. **(B)** ssDNA recombation enhancements for the strain lineags as measured via flow cytometry to measure the population fraction in which an sfGFP reporter could be turned off via incorporation of a permatuer stop codon. **(C)** The recombination effciency of BioDesignER compared to pTet-λ harboring modifications that interfere with mismatch repair (damOE,Δ*mutS*) (left, 37°C) and to canonical to p Tet-λ (control) to show retention of dsDNA recombination efficiency P-value from Mann-Whitney U-test; ns –not significnat. **(E)** Flow cytometry traces (top) with correspoding fold-change respponse curves (bottom) for each inducible, orthogonal regulator. Inducer conentrations used for flow cytome-try traces are: 0, 0.33, 0.67, 1.3, 3.3, 6.7, 33, 130 μM (arabinose); 0, 2, 5, 10, 20, 50, 100 μM (cumate); 0, 1, 2, 5, 10, 20, 50, 500 μM (IPTG).

### BioDesignER recombineering enhancements

#### ssDNA recombination enhancements

To quantify recombineering enhancements of key BioDesignER modifications, we measured ssDNA recombination rates for several strain intermediates. We integrated a dual fluorescent reporter cassette expressing both *sfGFP* and *mKate2* at a common genomic locus for each strain of the lineage and quantified ssDNA recombination efficiency. For each strain we transformed an oligo to inactivate *sfGFP* via incorporation of a premature stop codon. We also performed a control reaction in each case using water in place of oligo. After recovery and outgrowth we measured the fluorescence profiles of each strain using flow cytometry (Figure 2B). We observed increases in recombination efficiency at each modification stage with single cycle conversion rates improving from 1.6±0.1% (mean ± 1 standard deviation) in pTet-λ to 25.4±1.0% in BioDesignER.

To investigate the efficacy of mismatch repair evasion on recombination efficiency, we compared BioDesignER against pTet-λ derivative strains containing mismatch repair modifications and against two standard Δ*mutS* recombineering variants, EcNR2 and EcNR2.nuc5-. BioDesignER (25.4±1.0%) exhibits much higher recombination efficiency than pTet-λ with *dam* over-expression (damOE, 6.91±0.19%) or Δ*mutS* (12.9±1.7%) as hypermutagenesis strategies (Figure 2C, left panel). Performing the same recombineering experiments at 30°C and comparing to EcNR2 and EcNR2.nuc5-, which are constrained to growth at 30°C, we found that BioDesignER (13.6±1.2%) exhibited recombination rates comparable to EcNR2 (14.5±2.1%), yet approximately three-fold lower than EcNR2.nuc5- (37.7±3.8%) (Figure 2C, right panel). We were surprised to find that the recombineering efficiency of BioDesignER decreased by nearly two-fold when grown at a lower temperature.

#### dsDNA recombination enhancements

Knocking out endogenous exonucleases has been reported to significantly reduce or abolish dsDNA recombination efficiency (39). We measured the efficiency of dsDNA recombination in pTet-λ and BioDesignER and found no significant reduction in recombination efficiency (Figure 2D). This suggests that λ-*exo* is sufficient to process dsDNA recombination templates in the absence of multiple host exonucleases. A previous study reported that dsDNA recombination is at least an order of magnitude less efficient in a four-nuclease deficient genotype (Δ*exoX*, Δ*recJ*, Δ*xseA*, Δ*xonA*) with abolished dsDNA recombination activity in a three-nuclease (Δ*recJ*, Δ*xseA*, Δ*xonA*) knockout (39). We note here that we were successful in generating dsDNA recombinants in EcNR2.nuc5- at a similar efficiency to EcNR2 with no alteration to the recombineering protocol, suggesting that another nuclease is aiding dsDNA recombination in *E. coli* or that recombination can occur through an exonuclease-independent mechanism.

### Control of multiple independent regulators

BiodesignER expresses transcriptional regulators that utilize four independent small-molecule inducers to allow multi-input control of synthetic circuits, biosynthetic pathways, or gene editing tools. The strain produces the repressors TetR, LacI and CymR as well as the activator AraC. TetR is expressed from the λ prophage element native to EcNR1. We incorporated the transcriptional overexpression allele *lacI*^Q1^ to boost LacI production, which allows efficient regulation of multi-copy plasmids (40). We also introduced the tight and titratable regulator CymR (41), which is inactivated by the small molecule cumate. To improve gene regulation by arabinose we replaced the arabinose-sensitive promoter of the *araE* transporter gene with a constitutive promoter to eliminate all-or-none expression and allow titratable induction (42). In conjunction with this modification, we introduced a constitutive promoter to drive expression of AraC and deleted the *araBAD* operon to eliminate arabinose degradation via catabolism.

To characterize the induction profiles of each regulator, we quantified the fluorescence levels and growth rates of cells transformed with multi-copy plasmids. We constructed a set of GFP expression plasmids with promoters responsive to each regulator (Figure 2E, see Supplementary Figure S2 for sequence-level details) and transformed each plasmid into BioDesignER. Gene expression profiles were characterized by measuring single-cell fluorescence and bulk growth and fluorescence.

Fold-change induction for each regulator increased with plasmid copy number while no leaky expression was observed for low-copy and medium-copy plasmids. For plasmids with low-copy replication origin pSC101, we observed mean fold-change induction levels of 107, 68, and 20 for arabinose, cumate, and IPTG, respectively. For plasmids with medium-copy replication origin p15A, we observed mean fold change induction levels of 146, 184, and 30 for arabinose, cumate, and IPTG, respectively. In both copy-number contexts, GFP expression with no inducer was indistinguishable from a control plasmid lacking *gfp*. We found that repressor levels were insufficient to fully repress GFP expression on plasmids with the ColE1 replication origin. We note that AraC-regulated GFP expression saturates near 33 μM (5 μg/mL, 0.0005%) arabinose, a much lower saturation point than common plasmid-based systems (0.1% arabinose).

Single-cell distributions observed through flow cytometry revealed unimodal distributions of GFP expression for nearly all induction conditions (Figure 2E). GFP expression from both cumate- and IPTG-responsive promoters produced monotonic, decreasing coefficient of variation noise profiles for increasing inducer levels (Supplementary Figure S3). For arabinose induction, despite introducing modifications consistent with Khlebnikov *et al*., we observed significant cell-cell variability at two intermediate arabinose levels. Specifically, we observed a maximum coefficient of variation at 3.3 uM (Supplementary Figure S3), manifest in Figure 2E as the broad, weakly bimodal fluorescence distribution.

### Characterization of genomic integration Safe Sites

#### Genome Integration Safe Sites

To aid identifying genomic loci that provide reliable gene expression and recombination efficiency for future engineering efforts, we characterized a curated list of integration loci across the *E. coli* K-12 genome. The resulting eight genomic loci, termed Safe Sites, were chosen based on several criteria to minimize disruption to local chromosomal context upon integration of synthetic DNA constructs (Figure 3A, Supplementary Figure S4). Specifically, the integration Safe Sites are intergenic regions located between two convergently transcribed, non-essential genes that do not exhibit any phenotypes or growth defects across the majority of biochemical conditions screened in previous high-throughput studies (14, 43) (Supplementary Table S5), and contain no annotated features (small RNAs, promoters, transcription factor binding sites) according to RegulonDB (44) (Supplementary Table S6).

To characterize gene expression variation across the chromosome, we measured the expression of dual-fluorescent reporters (*sfGFP, mKate2*) integrated into BioDesignER at each Safe Site. We observed a linear decrease in expression for both sfGFP (pearson r_sfGFP,arm1_ = −0.91, pearson r_sfGFP,arm2_= −0.65, p_sfGFP_ < 0.05, permutation test) and mKate2 (pearson r_mKate,arm1_ = −0.85, pearson r_mKate,arm2_ = 0.51, p_mKate_ < 0.05, permutation test) reporters with respect to distance from the chromosomal origin (Figure 3B). This result was consistent with expected variations in local chromosomal copy number due to bi-directional replication dynamics during growth (45, 46). Interestingly, we observed a much stronger correlation of expression to distance from replication origin for chromosome Arm 1, though mKate2 expression at Safe Site 8 was a low outlier. We also assessed the effect of integration at each Safe Site on cellular fitness by measuring growth rates for each integration strain. We observed that, in general, genomic integration and expression from each Safe Site did not reduce growth rate, though Safe Site 8 displayed a nominal decrease when grown under aTc induction (Supplementary Figure S5). The two unexpected results at Safe Site 8 suggest that it may not be a reliable locus for integration.

**Figure 3.**
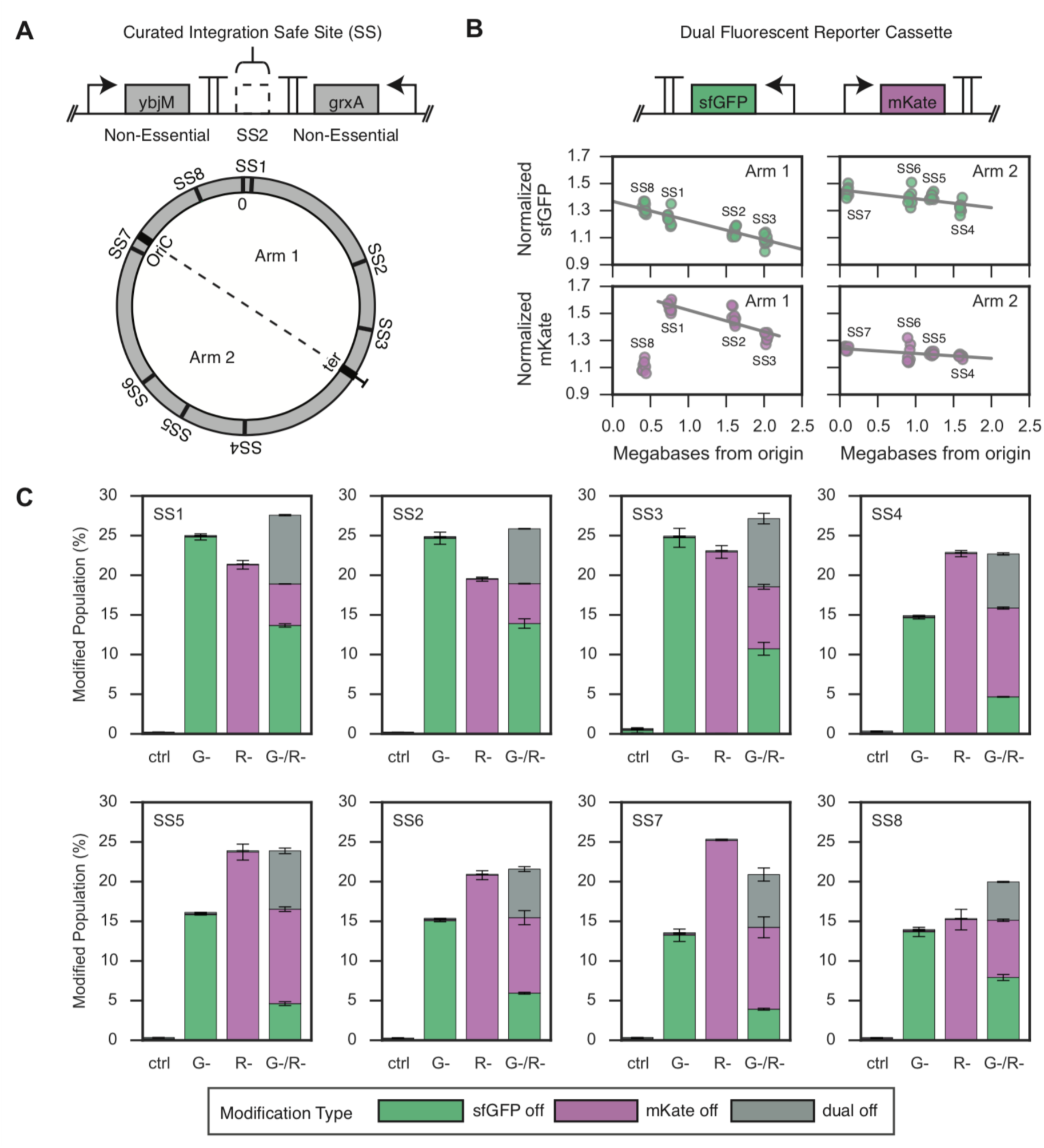
Recombination Characterization. **(A)** Circular map of the BioDesignER chromosome with Safe Sites mapped to corresponding genome position and chromosomal arm (replichore). **(B)** Genetic architecture of dual fluorescent reporter construct (top) and observed expression of reporters when integrated at each Safe Site on the chromosome (bottom). Replicate measurements of normalized expression levels for each reporter arrayed by chromosomal arm on which construct is integrated. **(C)** ssDNA recombination rates at each Safe Site for four independent recombineering reactions. X-axis denotes transformed oligo(s) (Gfor *sfGFP*, Rfor *mKate*) or ctrl (water). Bar height corresponds to the mean of two measurements and error bars represent span of data. Stacked barplots for each reaction represent popula tion fractions containing one of three possible modifications (*sfGFP* off, *mKate* off, or dual off when both reporters inactivated).

#### Recombination Rates across Safe Sites

Changes in local chromosomal structure may lead to unexpected fluctuations in recombination efficiency at various locations across the genome. To characterize recombination efficiency as a function of chromosomal locus for BioDesignER, we performed three independent ssDNA oligo-mediated recombination reactions for the panel of eight Safe Site strains. For each strain we independently transformed (1) an oligo to inactivate *sfGFP*, (2) an oligo to inactivate *mKate2*, or (3) an oligo cocktail to inactivate both reporters. We also performed a control reaction in each case using water in place of oligo. For Safe Sites that lie on opposite sides of the replication fork, we designed appropriate oligos to ensure recombination targeting the lagging strand. We found that recombination rates were consistently high across the chromosome with Safe Sites displaying single cycle, single site conversion rates of 17.0±6.70% and 19.7±5.7% for *sfGFP* and *mKate*2, respectively (Figure 3C). We also report single cycle, multiplex conversion rates of 7.5±4.4% for the *sfGFP*, 7.9±2.9% for the *mKate2*, and 6.3±2.3% for both reporters when transformed with the dual oligo cocktail.

### Analysis of transient hypermethylation effects on mutagenesis

To investigate the effect of BioDesignER modifications on global mutation rate, we developed a mutagenesis detection assay with a single nucleotide target. The mutagenesis cassette utilizes chloramphenicol acetyltransferase (*cat*) gene translationally fused to green fluorescence gene *mNeon* (Figure 4A). This strategy allows estimation of mutation rate without mutant fitness biases and secondsite suppressor mutations observed in traditional fluctuation analyses such as rifampicin resistance (47). Following integration of the mutagenesis cassette at Safe Site 1, we introduced a TAA stop codon at Lys19 of *cat* via a single nucleotide mutation. Only mutations that convert the stop codon to alternate codons generate chloramphenicol resistant, green fluorescent colonies, thus eliminating the possibility of suppressor mutations occurring in the fluctuation test.

**Figure 4.**
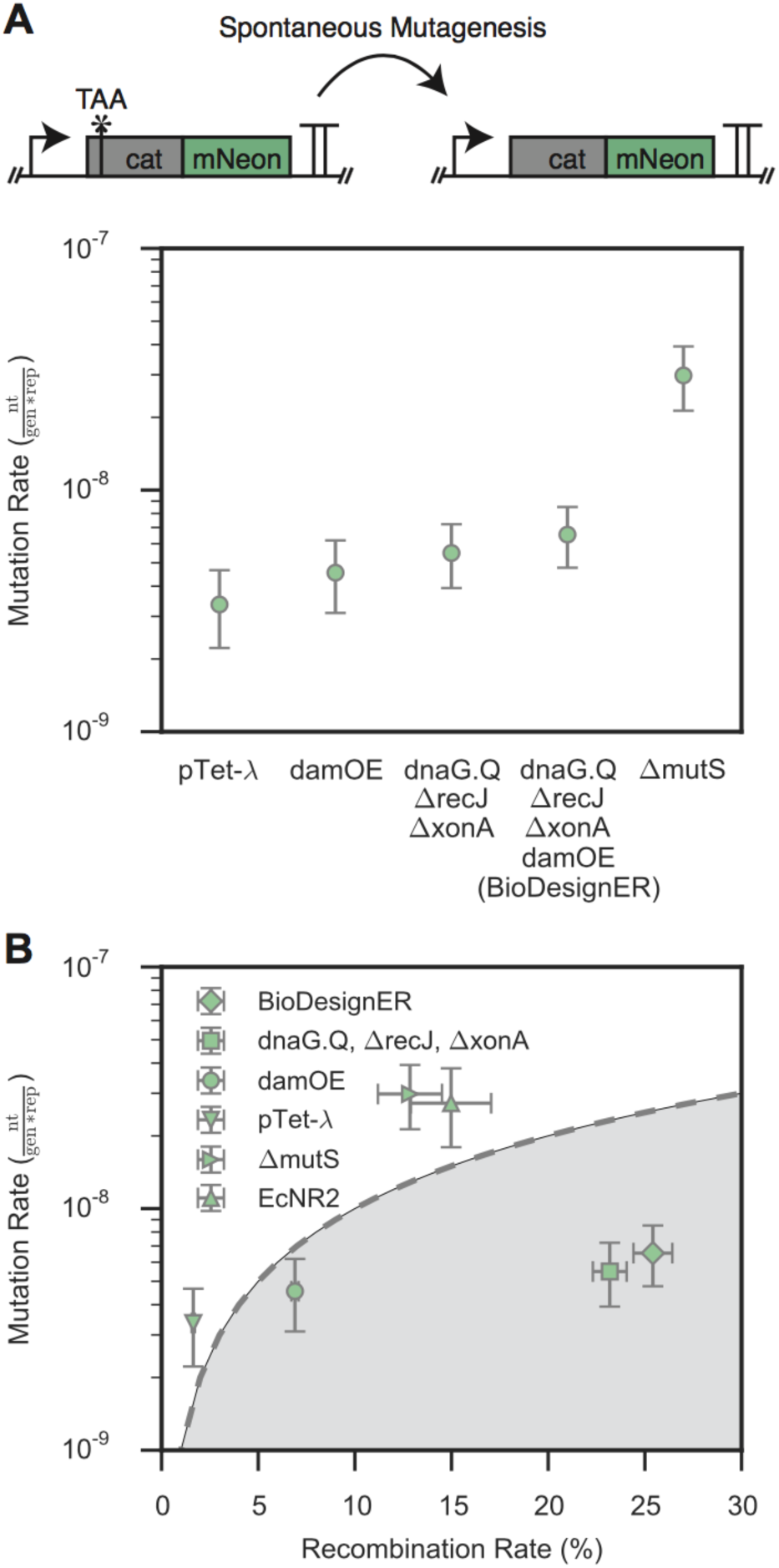
Mutaional Analysis. **(A)** Background mutation rates (nucleotides/genome/replication) as measured via a *cat-mNeon* fluctuation assay for various stages of BioDesignER strain contruction compared to an MMR-deficition (Δ*mutS*) strain derived from pTetλ Error bars represent 95% confidence intervals. **(B)** Single-cycle ssDNA recomination efficiency plotted against background mutation rate for each strain to show tradeoffs between recombination and mutation rates. The resulting tradeoff space represents the unit increase in mutation rate observed for a unit increase in recombination rate and is divide by y = β*x, where β = 10^-9^ is a characteristic scaling factor for the mutation rate X-error bars represent ± 1 standard deviation and Y-error bars represent 95% confidence intervals.

Using this assay, we benchmarked mutation rates of BioDesignER against (1) strains in the BioDesignER construction lineage, (2) EcNR2 (reference), and (3) MMR-deficient (control) strains pTet-λ *ΔmutS* and damOE. All assayed strains utilized the inactivated *cat-mNeon* cassette at Safe Site 1 as shown in Figure 4A. To allow EcNR2 to be compatible with the *cat-mNeon* fluctuation assay, we replaced the *cmR* selection cassette native to EcNR2 with *kanR*. Under comparable growth conditions, we estimated mutation rates of 3.36×10^-9^ (95% confidence interval (CI): 2.22-4.66×10^-9^) nucleotides/genome/replication, 4.55×10^-9^ (CI: 3.10-6.19×10^-9^), and 6.54×10^-9^ (CI: 4.77-8.51×10^-9^) for pTet-λ, damOE, and BioDesignER, respectively. By comparison, we observed mutation rates of 2.98×10^-8^ (CI: 2.13-3.93×10^-8^) for the control pTet-λ Δ*mutS*, which was similar to the rate of 2.73×10^-8^ (CI: 1.79-3.81×10^-8^) observed for EcNR2. For all strains assayed, all chloramphenicol-resistant colonies were also fluorescent. From a set of 54 individually sequenced chloramphenicol-resistant clones, we observed 8 unique genotypes arising from spontaneous mutations (Supplementary Figure S6).

To investigate the effect of λ-Red induction on global mutation rates and compare the mutagenic effect of dam over-expression to deletion of *mutS*, we tested the mutation rates for pTet-λ, damOE, and pTetλ *ΔmutS* both with and without aTc induction (Supplementary Figure S6). We found no effect on global mutation rates due to aTc induction (i.e. expression of the λ-Red machinery) in pTet-λ and pTet-λ *ΔmutS*. Consistent with prior work (31), we observed an increase in mutation rate for damOE under aTc induction - specifically, 2.4-fold in this work. Finally, we noted that even with aTc induction damOE was still less mutagenic than pTet-λ *ΔmutS*, suggesting that BioDesignER uniquely strikes a balance between on-target and off-target mutagenesis rates.

To quantify this balance, we compared the recombination and mutagenesis rates for a selection of control strains and BioDesignER (Figure 4B). The resulting trade-off space can be divided into two regimes where strains falling in the shaded region exhibit a favorable trade-off between recombination rate and mutation rate. BioDesignER falls in the favorable subspace, while MMR-deficient strains such as EcNR2 and the pTet-λ Δ*mutS* fall in the unfavorable regime above the tradeoff line. To summarize this result we introduce the metric recombineering fidelity, which we define as the product of fold-increase in recombination rate and fold-decrease in mutagenesis rate, each relative to EcNR2. Using this metric we calculate that BioDesignER exhibits 7.3-fold greater recombineering fidelity than EcNR2 (1.75-fold improvement in recombination rate and 4.17-fold decrease in mutagenesis rate) (Table 2).

**Table 2.** Comparison of recombineering fidelity factors for relevant strains.

| Strain | Recombination Efficiency | Background Mutation Rate | Recombineering Fidelity |
| --- | --- | --- | --- |
| pTet-λ | 1.6±0.1% | 3.36 × 10<sup>-9</sup> | 0.9 |
| EcNR2 | 14.5±2.1% | 2.73 × 10<sup>-8</sup> | 1.0 |
| BioDesignER | 25.4±1.0% | 6.54 × 10<sup>-9</sup> | 7.3 |

**Note:** EcNR2 data collected at 30°C, pTet-λ and BioDesignER at 37°C

### Multi-cycle recombineering rate enhancements

High single-cycle editing efficiency enables the rapid generation of genotypically diverse populations using multiplexed, cyclical recombineering workflows. To assess how well BioDesignER could generate a population with multiplex edits, we transformed a starting population with an oligo cocktail targeting multiple sites and tracked phenotypic diversity as a function of recombineering cycle for multiple strains. Specifically, we transformed BioDesignER harboring the *sfGFP-mKate2* fluorescence cassette with oligos to inactivate both reporters over four sequential recombineering cycles. In parallel, we compared BioDesigner to pTet-λ (Figure 5A), EcNR2, and EcNR2.nuc5- (Supplementary Figure S7) transformed with the same cocktail. BioDesignER exhibited high multiplex editing efficiency with nearly 60 percent of the population incorporating both edits (58.8±3.5%) by the fourth recombineering cycle (Figure 5A), thus outperforming EcNR2 (15.9±3.0%) and showing similar efficiency to EcNR2.nuc5- (54.3±5.6%) (Figure 5B).

**Figure 5.**
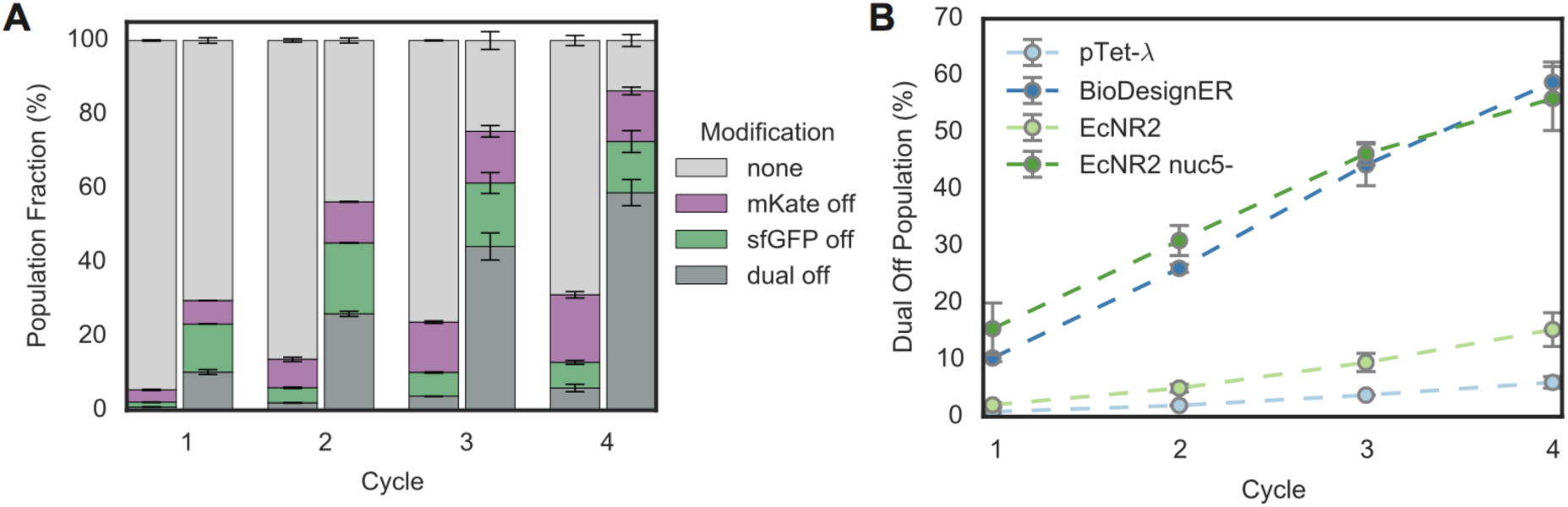
Multi-Cycle Recombineering. **(A)** The fraction of eah genotype (i.e. modification type) was measured via flow cytometry for pTetλ (left) and BioDesignER (right) after each cycle of recombineering. Errors bars represent ± 1 standard deviation. **(B)** The fraction of each strain population in which boh markers werwe edited (dual off genotype) is shown across all four recombineering cycles. Error bars represent α 1 standard deviation.

Given the higher single-cycle conversion rate of EcNR2.nuc5- compared to BioDesignER (Figure 2C, right panel), we were surprised by the comparable performance of the two strains over multiple recombineering cycles. We partly attribute this parity to uncharacteristically low and sporadic single-cycle efficiencies that we repeatedly observed for EcNR2.nuc5- replicates (Supplementary Figure S7). Regardless, while both BioDesignER and EcNR2.nuc5- exhibited similar multiplex editing efficiencies, EcNR2.nuc5- requires culturing at 30-32°C and is a persistent mutator, which increases recombineering cycle time and basal mutation rate, respectively thus limiting its overall utility as a reliable strain for multiplex genome editing.

## DISCUSSION

High-efficiency genome engineering in bacteria enables breadth (24) and depth (17) explorations of genotypic diversity to enhance engineered behaviors. However, to date, no platform strain exists that incorporates a suite of core functions to provide efficient recombineering and regulate both genome engineering functions and cellular programs. BioDesignER is a high recombineering fidelity recombineering strain constructed to rapidly explore and optimize engineered functions. It incorporates many genomic modifications that increase recombination efficiency and reduce cycle time for recombineering workflows while minimizing off-target mutations. BioDesignER includes four independent inducible regulators to control recombineering and accommodate additional user designs. We have characterized eight Safe Site integration loci distributed across the genome and found that seven enable reliable gene expression and mutagenesis.

BioDesignER enables rapid selection-based recombineering workflows with no requirements for plasmid transformation or curing. Reliable engineering of sequential genome integrations with established recombineering approaches, such as the use of plasmids pSIM5 (12) or pKD20 (10), require transformation and curing procedures of plasmid-encoded recombineering functions for each integration stage. These requirements increase the time required for individual genome editing steps by multiple days. Anecdotally, we have found plasmid-based recombineering systems unreliable for conducting multiple editing cycles from a single transformation of the recombineering plasmid. We speculate that the failure to achieve multi-cycle genome editing from plasmid-based recombineering solutions may be related to the accumulation of mutations spurred by maintenance or leaky expression of λ-Red genes over many generations. In contrast, we have completed all of the scar-free DNA recombineering workflows reported here with no restoration or replacement of the minimized pTet λRed cassette.

We have increased recombineering fidelity in BioDesignER by striking a balance between recombination efficiency and mutagenesis rates. A high recombineering fidelity platform such as this may provide new avenues to multiplex genome remodeling using CRISPR-Cas9 techniques. CRISPR-Cas9 genome editing approaches in bacteria are limited by recombination efficiency to rescue doublestrand breaks. Linking CRISPR-Cas9 counterselection of native sequences with high-efficiency, multisite recombineering may allow concurrent selection of many modifications from a large bacterial population with little off-target activity, thereby enabling researchers to explore unprecedented genetic diversity.

While BioDesignER exhibits robust functionalities with respect to recombineering fidelity, comparing the recombination efficiency of the BioDesignER lineage to EcNR2-derived strains reveals inconsistent results related to culture temperatures. Specifically, we found a nearly two-fold reduction in recombination efficiency for BioDesignER at 30°C compared to 37°C, resulting in recombineering efficiencies similar to EcNR2 (Figure 2C). This reduction suggests some uncharacterized temperature-specific reduction in recombination efficiency and could reflect reduced ssDNA access to the replication fork, lower ssDNA half-life at reduced temperatures, or perhaps uncharacterized temperature-dependent expression of the λ-Red machinery from pTet.

While constructing BioDesignER, we developed multiple selection/counter-selection strategies that may be of general use for bacterial genome engineering. These strategies combine selection/counterselection and fluorescence screening components to accelerate scar-free genome engineering. Specifically, the genetic cassettes utilize selection/counter-selection of *thyA*, building on work from FRUIT (48). This approach requires two recombineering transformations: a dsDNA integration of the fluorescence-coupled *thyA* cassette at the target genomic locus followed by removal of the cassette using ssDNA or dsDNA. The genetic modification of interest can be incorporated at either integration stage. In comparison, CRISPR-based genome editing workflows, which are gaining popularity, require multiple steps including guide plasmid construction, co-transformation with Cas9, and subsequent curing. Thus, the selection/counter selection methodologies developed here allow a simple and effective approach to genome engineering.

Development of BioDesignER points to genome design strategies for next-generation biotechnology hosts. As synthetic biology matures, the application space is expanding beyond prototypical genetic circuits and metabolic pathways in laboratory environments to robust engineered functions in ecologies with high biotic and abiotic complexity, including soil, wastewater, and the human gut (49). Efficient and sustained activity of engineered functions in these environments will require programmed behaviors to be optimized in phylogenetically diverse microbes. We anticipate the integrative approach used to develop and characterize BioDesignER can be used as a template to develop high-efficiency recombineering platforms for new bacterial hosts.

## ACKNOWLEDGEMENTS

The authors would like to thank Vivek Mutalik for providing the *cymR* expression construct and the core promoter sequence used to design the Tet-based λ-Red expression system. We would also like to thank the Li Ka Shing Flow Cytometry Facility at UC Berkeley.

## FUNDING

This work was supported by the Department of Energy Genome Science program within the Office of Biological and Environmental Research [grant number DE-SC008812, Funding Opportunity Announcement DE-FOA-0000640]. H.S.R is supported by a National Science Foundation (NSF) Graduate Research Fellowship and a National Institutes of Health (NIH) Genomics and Computational Biology Training Program [5T32HG000047-18]. Funding for open access charge: DE-SC008812.

### Conflict of interest statement

None declared.

## REFERENCES

1. Khalil, A.S. and Collins, J.J. (2010) Synthetic biology: applications come of age. Nat Rev Genet, 11, 367–379.

2. Weber, W. and Fussenegger, M. (2011) Emerging biomedical applications of synthetic biology. Nat Rev Genet, 13, 21–35.

3. Tyo, K.E.J., Ajikumar, P.K. and Stephanopoulos, G. (2009) Stabilized gene duplication enables long-term selection-free heterologous pathway expression. Nat Biotechnol, 27, 760–765.

4. Friehs, K. (2004) Plasmid copy number and plasmid stability. Adv Biochem Eng Biotechnol, 86, 47–82.

5. Bassalo, M.C., Garst, A.D., Halweg-Edwards, A.L., Grau, W.C., Domaille, D.W., Mutalik, V.K., Arkin, A.P. and Gill, R.T. (2016) Rapid and Efficient One-Step Metabolic Pathway Integration in E. coli. ACS Synth. Biol., 5, 561–568.

6. Lee, J.W., Gyorgy, A., Cameron, D.E., Pyenson, N., Choi, K.R., Way, J.C., Silver, P.A., Del Vecchio, D. and Collins, J.J. (2016) Creating Single-Copy Genetic Circuits. Molecular Cell, 63, 329–336.

7. Esvelt, K.M. and Wang, H.H. (2013) Genome-scale engineering for systems and synthetic biology. Molecular Systems Biology, 9, 641–641.

8. Bryant, J.A., Sellars, L.E., Busby, S.J.W. and Lee, D.J. (2014) Chromosome position effects on gene expression in Escherichia coli K-1Nucleic Acids Res, 42, 11383–11392.

9. Murphy, K.C. (1998) Use of bacteriophage lambda recombination functions to promote gene replacement in Escherichia coli. J. Bacteriol., 180, 2063–2071.

10. Datsenko, K.A. and Wanner, B.L. (2000) One-step inactivation of chromosomal genes in Escherichia coli K-12 using PCR products. Proc Natl Acad Sci USA, 97, 6640–6645.

11. Ellis, H.M., Yu, D., DiTizio, T. and Court, D.L. (2001) High efficiency mutagenesis, repair, and engineering of chromosomal DNA using single-stranded oligonucleotides. Proc Natl Acad Sci USA, 98, 6742–6746.

12. Datta, S., Costantino, N. and Court, D.L. (2006) A set of recombineering plasmids for gram-negative bacteria. Gene, 379, 109–115.

13. Sharan, S.K., Thomason, L.C., Kuznetsov, S.G. and Court, D.L. (2009) Recombineering: a homologous recombination-based method of genetic engineering. Nature Protocols, 4, 206–223.

14. Baba, T., Ara, T., Hasegawa, M., Takai, Y., Okumura, Y., Baba, M., Datsenko, K.A., Tomita, M., Wanner, B.L. and Mori, H. (2006) Construction of Escherichia coli K-12 in-frame, single-gene knockout mutants: the Keio collection. Molecular Systems Biology, 2, 1–11.

15. Warner, J.R., Reeder, P.J., Karimpour-Fard, A., Woodruff, L.B.A. and Gill, R.T. (2010) Rapid profiling of a microbial genome using mixtures of barcoded oligonucleotides. Nat Biotechnol, 28, 856–862.

16. Freed, E.F., Winkler, J.D., Weiss, S.J., Garst, A.D., Mutalik, V.K., Arkin, A.P., Knight, R. and Gill, R.T. (2015) Genome-Wide Tuning of Protein Expression Levels to Rapidly Engineer Microbial Traits. ACS Synth. Biol., 4, 1244–1253.

17. Garst, A.D., Bassalo, M.C., Pines, G., Lynch, S.A., Halweg-Edwards, A.L., Liu, R., Liang, L., Wang, Z., Zeitoun, R., Alexander, W.G., et al. (2016) Genome-wide mapping of mutations at single-nucleotide resolution for protein, metabolic and genome engineering. Nat Biotechnol, 35, 1–12.

18. Wang, H.H., Isaacs, F.J., Carr, P.A., Sun, Z.Z., Xu, G., Forest, C.R. and Church, G.M. (2009) Programming cells by multiplex genome engineering and accelerated evolution. Nature, 460, 894–898.

19. Isaacs, F.J., Carr, P.A., Wang, H.H. and Lajoie, M.J. (2011) Precise manipulation of chromosomes in vivo enables genome-wide codon replacement. Science, 10.1126/science.1204763.

20. Lajoie, M.J., Rovner, A.J., Goodman, D.B., Aerni, H.-R., Haimovich, A.D., Kuznetsov, G., Mercer, J.A., Wang, H.H., Carr, P.A., Mosberg, J.A., et al. (2013) Genomically recoded organisms expand biological functions. Science, 342, 357–360.

21. Lajoie, M.J., Kosuri, S., Mosberg, J.A., Gregg, C.J., Zhang, D. and Church, G.M. (2013) Probing the limits of genetic recoding in essential genes. Science, 342, 361–363.

22. Wang, H.H., Kim, H., Cong, L., Jeong, J. and Bang, D. (2012) Genome-scale promoter engineering by coselection MAGE. Nature, 10.1038/nmeth.1971.

23. Wang, H.H., Huang, P.-Y., Xu, G., Haas, W., Marblestone, A., Li, J., Gygi, S.P., Forster, A.C., Jewett, M.C. and Church, G.M. (2012) Multiplexed in vivo His-tagging of enzyme pathways for in vitro single-pot multienzyme catalysis. ACS Synth. Biol., 1, 43–52.

24. Zeitoun, R.I., Garst, A.D., Degen, G.D., Pines, G., Mansell, T.J., Glebes, T.Y., Boyle, N.R. and Gill, R.T. (2015) Multiplexed tracking of combinatorial genomic mutations in engineered cell populations. Nat Biotechnol, 33, 1–10.

25. Zeitoun, R.I., Pines, G., Grau, W.C. and Gill, R.T. (2017) Quantitative Tracking of Combinatorially Engineered Populations with Multiplexed Binary Assemblies. ACS Synth. Biol., 6, 619–627.

26. Glickman, B.W. and Radman, M. (1980) Escherichia coli mutator mutants deficient in methylation-instructed DNA mismatch correction. Proc Natl Acad Sci USA, 77, 1063–1067.

27. Schaaper, R.M. and Dunn, R.L. (1987) Spectra of spontaneous mutations in Escherichia coli strains defective in mismatch correction: the nature of in vivo DNA replication errors. Proc Natl Acad Sci USA, 84, 6220–6224.

28. Sawitzke, J.A., Thomason, L.C., Costantino, N., Bubunenko, M., Datta, S. and Court, D.L. (2007) Recombineering: in vivo genetic engineering in E. coli, S. enterica, and beyond. Meth. Enzymol., 421, 171–199.

29. Wang, H.H., Xu, G., Vonner, A.J. and Church, G. (2011) Modified bases enable high-efficiency oligonucleotide-mediated allelic replacement via mismatch repair evasion. Nucleic Acids Res, 39, 7336–7347.

30. Nyerges, Á., Csörgő, B., Nagy, I., Latinovics, D., Szamecz, B., Pósfai, G. and Pál, C. (2014) Conditional DNA repair mutants enable highly precise genome engineering. Nucleic Acids Res, 42, e62–e62.

31. Lennen, R.M., Nilsson Wallin, A.I., Pedersen, M., Bonde, M., Luo, H., Herrgård, M.J. and Sommer, M.O.A. (2016) Transient overexpression of DNA adenine methylase enables efficient and mobile genome engineering with reduced off-target effects. Nucleic Acids Res, 44, e36–e36.

32. Sarkar, S., Ma, W.T. and Sandri, G.H. (1992) On fluctuation analysis: a new, simple and efficient method for computing the expected number of mutants. Genetica, 85, 173–179.

33. Ma, W.T., Sandri, G.H. and Sarkar, S. (1992) Analysis of the Luria–Delbrück distribution using discrete convolution powers. Journal of Applied Probability.

34. Sergueev, K., Yu, D., Austin, S. and Court, D. (2001) Cell toxicity caused by products of the p(L) operon of bacteriophage lambda. Gene, 272, 227–235.

35. Lawther, R.P., Calhoun, D.H., Adams, C.W., Hauser, C.A., Gray, J. and Hatfield, G.W. (1981) Molecular basis of valine resistance in Escherichia coli K-1 Proc Natl Acad Sci USA, 78, 922–925.

36. Lawther, R.P., Calhoun, D.H., Gray, J., Adams, C.W., Hauser, C.A. and Hatfield, G.W. (1982) DNA sequence fine-structure analysis of ilvG (IlvG+) mutations of Escherichia coli K-1J. Bacteriol., 149, 294–298.

37. Tedin, K. and Norel, F. (2001) Comparison of ?relA strains of Escherichia coli and Salmonella enterica serovar Typhimurium suggests a role for ppGpp in attenuation regulation of branched-chain …. J. Bacteriol., 10.1128/JB.183.21.6184-6196.2001.

38. Lajoie, M.J., Gregg, C.J., Mosberg, J.A., Washington, G.C. and Church, G.M. (2012) Manipulating replisome dynamics to enhance lambda Red-mediated multiplex genome engineering. Nucleic Acids Res, 40, e170–e170.

39. Mosberg, J.A., Gregg, C.J., Lajoie, M.J., Wang, H.H. and Church, G.M. (2012) Improving lambda red genome engineering in Escherichia coli via rational removal of endogenous nucleases. PLoS ONE, 7, e44638.

40. Glascock, C.B. and Weickert, M.J. (1998) Using chromosomal lacIQ1 to control expression of genes on high-copy-number plasmids in Escherichia coli. Gene, 223, 221–231.

41. Choi, Y.J., Morel, L., Le François, T., Bourque, D., Bourget, L., Groleau, D., Massie, B. and Míguez, C.B. (2010) Novel, versatile, and tightly regulated expression system for Escherichia coli strains. Appl. Environ. Microbiol., 76, 5058–5066.

42. Khlebnikov, A., Skaug, T. and Keasling, J.D. (2002) Modulation of gene expression from the arabinose-inducible araBAD promoter. J Ind Microbiol Biotechnol, 29, 34–37.

43. Price, M.N., Wetmore, K.M., Waters, R.J., Callaghan, M., Ray, J., Liu, H., Kuehl, J.V., Melnyk, R.A., Lamson, J.S., Suh, Y., et al. (2018) Mutant phenotypes for thousands of bacterial genes of unknown function. Nature, 557, 503–509.

44. Gama-Castro, S., Salgado, H., Santos-Zavaleta, A., Ledezma-Tejeida, D., Muñiz-Rascado, L., García-Sotelo, J.S., Alquicira-Hernández, K., Martínez-Flores, I., Pannier, L., Castro-Mondragón, J.A., et al. (2016) RegulonDB version 9.0: high-level integration of gene regulation, coexpression, motif clustering and beyond. Nucleic Acids Res, 44, D133–43.

45. Bipatnath, M., Dennis, P.P. and Bremer, H. (1998) Initiation and velocity of chromosome replication in Escherichia coli B/r and K-1 J. Bacteriol., 180, 265–273.

46. Reynolds, T.S. and Gill, R.T. (2015) Quantifying Impact of Chromosome Copy Number on Recombination in Escherichia coli. ACS Synth. Biol., 4, 776–780.

47. Lee, H., Popodi, E., Tang, H. and Foster, P.L. (2012) Rate and molecular spectrum of spontaneous mutations in the bacterium Escherichia coli as determined by whole-genome sequencing. Proc. Natl. Acad. Sci. U.S.A., 109, E 2774–83.

48. Stringer, A.M., Singh, N., Yermakova, A., Petrone, B.L., Amarasinghe, J.J., Reyes-Diaz, L., Mantis, N.J. and Wade, J.T. (2012) FRUIT, a scar-free system for targeted chromosomal mutagenesis, epitope tagging, and promoter replacement in Escherichia coli and Salmonella enterica. PLoS ONE, 7, e44841.

49. Venturelli, O.S., Egbert, R.G. and Arkin, A.P. (2016) Towards Engineering Biological Systems in a Broader Context. J. Mol. Biol., 428, 928–944.

